# Filopodial Mechanotransduction is regulated by Angiotensin-Converting Enzyme 2 (ACE2) and by SARS-CoV-2 spike protein

**DOI:** 10.1101/2024.03.01.581813

**Authors:** Wei He, Chien-Ting Wu, Peter K. Jackson

**Affiliations:** Baxter Laboratory, Department of Microbiology & Immunology and Department of Pathology, Stanford University School of Medicine, Stanford, CA 94305, USA

## Abstract

Filopodia are dynamic, actin-rich cellular protrusions, increasingly linked to cellular mechanotransduction. However, how dynamic filopodia translate external mechanical cues remains poorly understood. Recent studies show that the SARS-CoV-2 spike (S) protein binds the ACE2 receptor on airway multicilia and that cilia are required for viral infection(*1*) and sufficient to induce filopodial extension and viral binding. To test if spike protein is sufficient to induce filopodial expansion, we employed live-cell single-particle tracking with quantum dots targeting ACE2, to reveal a robust filopodia extension and virus binding mechanism requiring the enzymatic activity of ACE2. Using time-lapse imaging, we reveal that spike protein binding to filopodia is associated with intracellular actin remodeling, alterations in bulk cell stiffness, and an elevation in intracellular calcium levels linked to actin-rearrangement, filopodia initiation, and persistence. We propose the activation of ACE2 creates an active signaling and mechanosensory environment within adherent cells and airway epithelial cells that allows the remodeling of actin in filopodia to trap virus and potentially organize viral exit from cells. [164 words]

**One-Sentence Summary:** SARS-CoV-2 spike protein activates calcium and actin dynamics to enable filopodial extension and virus binding

## Main Text

Filopodia are dynamic cellular structures pivotal in the intricate process of sensing the surrounding microenvironment(*2*). Their critical roles are exemplified in large scale cell migration in tissue morphogenesis instances such as axon pathfinding(*3*), embryonic dorsal closure(*4-6*), and endothelial cell (EC) angiogenesis(*7-9*). While the presence and activity of filopodia are widely recognized as important indicators of cell responsiveness to external cues, the process by which cells sense and transduce signals from filopodia remains mostly elusive. The inadequate understanding of the role of filopodia is partly due to the lack of a robust and versatile filopodia model system. Due to the dynamic and spontaneous nature of filopodia, it is difficult to correlate intracellular events when filopodia formation is infrequent (as in the case of growth factor-coated beads), of short duration (as in the case of starvation or cell migration) or lacks clear initiation (as in most resting tissues).

Our recent findings demonstrate that viral infection of airway organoids show strong extension and branching of filopodia at later time points (24-48 hours). Viral particles avidly attach to these new filopodia suggesting that filopodia may be important for regulating viral exit from the airway. The mechanism from SARS-CoV-2 infection to filopodial expansion is supported by phosphoproteomics data showing viral stimulation of phosphorylation of filopodia proteins like ezrin and PAK1 or PAK4 kinases.

To investigate signaling the crosstalk between ACE2 enzymatic activity and ACE2 signals induced by the SARS-CoV-2 Spike receptor-binding domain (S-RBD), we employed live-cell single-particle tracking techniques to monitor ACE2 labeled with quantum dots. Our findings revealed that the ACE2-S-RBD interaction surprisingly manifests as an ideal cell model for studying filopodia, offering controlled initiation, abundant filopodia formation, and an extended observation window. This innovative filopodia cell model has, for the first time, provided insights to model how cells mechanically respond to persistent and substantial filopodia interactions over time.

Upregulation of the mechanosensitive channel piezo1 is associated with pulmonary hypertension(*10, 11*), the most prevalent cardiovascular comorbidity in patients infected with coronavirus disease 2019 (COVID-19)(*12*), and with post COVID-19 pulmonary fibrosis(*13, 14*). Only recently, a study confirmed that treatment with S-RBD is sufficient to induce upregulation of several mechanosensitive channels, including Piezo1, leading to prolonged intracellular free calcium levels in pulmonary vascular endothelial cells (*15*). The mechanism through which mechanosensation is realized via the S-RBD-ACE2 interaction remains unknown. Combining our working hypothesis of the filopodia induction by the S-RBD-ACE2 interaction, (*1*), we tested the filopodia induction model as a readout in motile cilia beating frequency (CBF) in airway epithelial cells. As a result, a new perspective on ACE2’s function as a mechanotransduction activator of filopodia emerges in both adherent cells and airway epithelial cells.

## Quantum dot labeling of ACE2 revealed its initiation and accumulation on filopodia

The multifaceted roles of ACE2 extend predate its function as a receptor for the SARS-CoV-2 virus. A primary focus of research has been directed towards its involvement in catalyzing angiotensin II (Ang II) within the classical renin–angiotensin system (RAS) pathway, a pivotal regulatory mechanism governing blood pressure. Notably, the binding sites of the virus and Ang II on ACE2 are observed to be non-competitive binding sites. Ang II binds as a substrate in the catalytic pocket of ACE2, whereas the S-RBD recognizes the distant N-terminal region of ACE2.

Despite this structural distinction, clinical observations in COVID-19 patients reveal intriguing parallels between SARS-CoV-2 infection and the upregulation of Ang II. Common outcomes include hypertension (*12*) and increased autoantibodies targeting Ang II (*16*). To elucidate potential crosslinks in ACE2 signaling triggered by Ang II or S-RBD, we employed quantum dots (QDs) to label ACE2, and live cell single-particle tracking was conducted on A549 cells featuring a knock-in ACE2 expressing human ACE2 (A549 ACE2 cells).

We first used an anti-ACE2 antibody to label ACE2 protein by attaching a QD to the antibody (anti-ACE2-QD, Fig. 1A). This strategy offers the advantage of capturing the pre-activated state of ACE2. In this state, ACE2 trajectories exhibit diffusion at a velocity comparable to other transmembrane proteins (*17*). After treatment with Ang II or S-RBD, a notable 3-8-fold reduction in the diffusion speed of anti-ACE2-QD was observed (Movie S1). Furthermore, a concurrent enhancement of the sizes of QDs was noted (Fig. 1B, C), implying potential clustering or endocytosis.

**Fig. 1.**
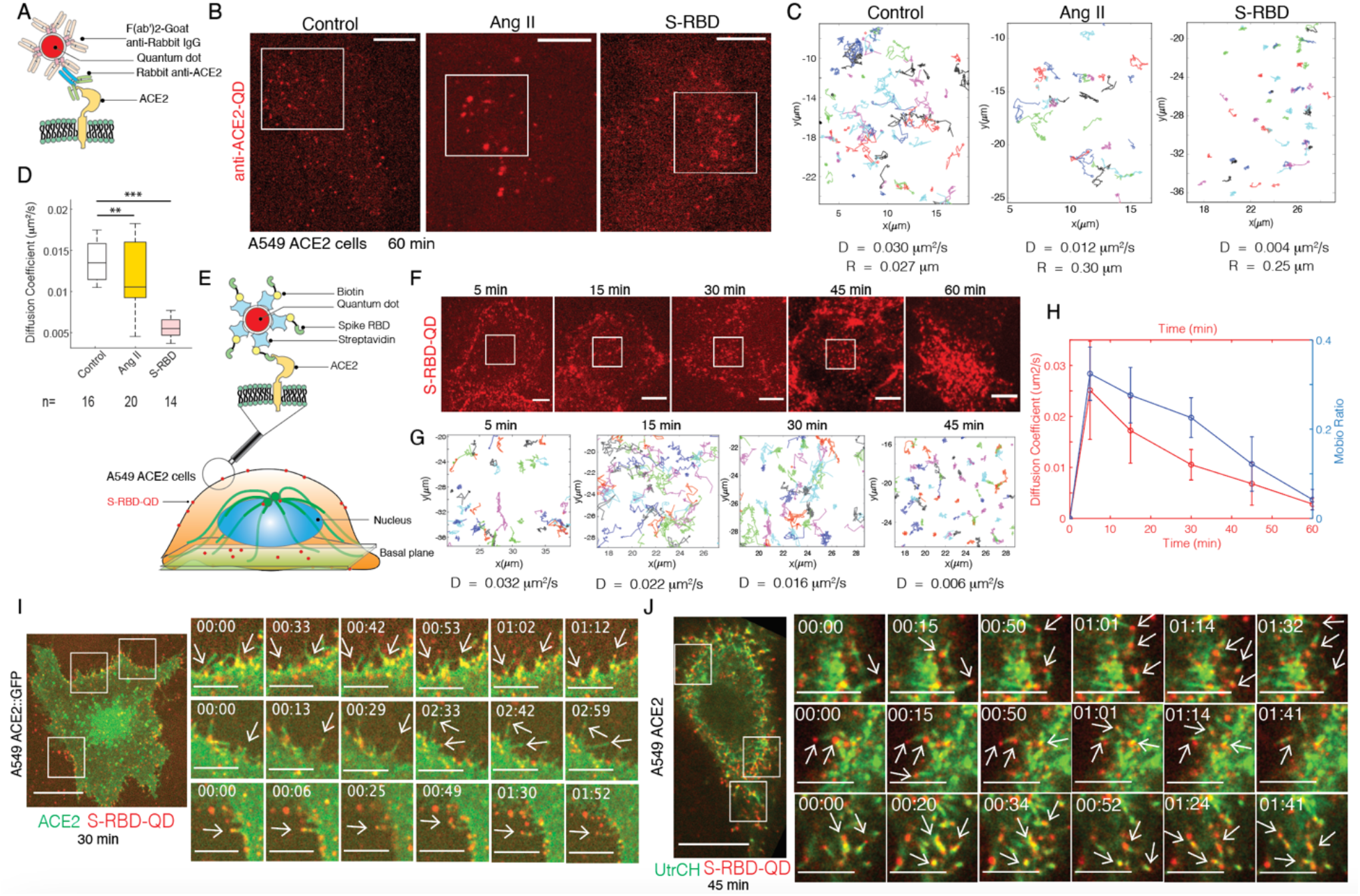
Quantum dot labeling of ACE2 revealed its initiation and accumulation on filopodia. **(A)** Illustration of Quantum Dot (QD) Labeling of ACE2 by Anti-ACE2 Antibody: Rabbit anti-ACE2 primary antibody (green and blue) recognizes the first 50 amino acids of the transmembrane ACE2 (yellow). Quantum dot (red) binds through the F(ab’) fragment (pink) of the secondary antibody coated on the QD (red). **(B)** Representative Images of Anti-ACE2-QD (Red): Obtained from live A549 ACE2 cells after 60 minutes of treatment from Control (no treatment), Ang II (200nM), and S-RBD (100nM) treatment. Scale bar is 10 μm. **(C)** Tracking Trajectories: In the duration of 3 minutes of anti-ACE2-QD from three example cells. Crop regions are as shown in the white box regions from (B) corresponding to Control, Ang II, and S-RBD treatment. The trajectories are labeled in random colors. Diffusion coefficients and the average QD sizes calculated from the cropped region are presented under each trajectory plot. **(D)** Diffusion Coefficients Box Plot: Calculated for anti-ACE2-QD obtained in a 3-minute duration from Control, Ang II (200nM), and S-RBD (100nM) treatment conditions. (\*\**P* <0.01; \*\*\**P* <0.001). Cell numbers (n) for each calculation is present under each box. **(E)** Illustration of QD Labeling of ACE2 by S-RBD: S-RBD (green), biotinylated, recognizes the N-terminal of the transmembrane ACE2 (yellow). Quantum dot (red), conjugated with streptavidin, binds to biotin. **(F)** Representative Time-Lapse Images of S-RBD-QD (Red): Obtained at the basal plane (as illustrated in (E)) from live A549 ACE2 cells after S-RBD-QD treatment for 5, 15, 30, 45, 60 minutes. Scale bar is 10 μm. **(G)** Tracking Trajectories: In the duration of 2 minutes of S-RBD-QD from four example cells. Crop regions are as shown in the white box regions from (F) corresponding to 5, 15, 30, 45 minutes. Diffusion coefficients calculated from the cropped region are presented under each trajectory plot. **(H)** Diffusion Coefficient (Red) and Mobile Ratio (Blue) Plots: In a time-lapse of 5, 15, 30, 45, 60 minutes after S-RBD-QD treatment (S-RBD 100nM) on A549 ACE2 cells at the basal plane. **(I)** Time-Lapse Images of S-RBD-QD (Red) on A549 ACE2::GFP Cells: Demonstrating S-RBD-QDs staying at the tip of spouting filopodia. Three random crop regions from the whole cell (S-RBD treatment 30 mins) on the left (Scale bar 10 μm) are amplified on the right (Scale bar 5 μm) with white arrows pointing to the growing tips. **(J)** Time-Lapse Images of S-RBD-QD (Red) on an A549 ACE2 Cell: (S-RBD treatment 45 mins) Transiently transfected with UtrCH (green), demonstrating S-RBD-QDs staying at existing filopodia. Three random crop regions from the whole cell view on the left (Scale bar 10 μm) are amplified on the right (Scale bar 5 μm) with white arrows pointing to the tips of filopodia.

RBD-conjugated QD has been previously reported to effectively induce endocytosis of ACE2, and slowdown of QD motility is regarded as reflection of intracellular confinement(*18*). The specificity of biotinylated S-RBD, which binds to streptavidin-conjugated QD (S-RBD-QD), to ACE2 is confirmed by the reduced labeling of QD when co-treated with soluble ACE2 (sACE2) or by observing entirely negative labeling in WT A549 cells lacking expressed ACE2 receptor (Fig. S1). To further elucidate the cause and cellular localization of ACE2 reduced motility, we applied S-RBD-QD to achieve faster and denser labeling of ACE2, enabling time-lapse characterization of the diffusion coefficient and mobile ratio of S-RBD-QD-bound ACE2 from 5 minutes to 60 minutes. To our surprise, an additional population of ACE2 becomes immobile on the cell surface, in both a time-dependent (Fig. 1F) and dose-dependent manner. Both alterations are sensitive to the ACE2 inhibitor (Fig. S2). At the 5 min time point, the diffusion speed of S-RBD-QD is at a similar level as untreated anti-ACE2-QD, while after 60 min the diffusion speed to that from anti-ACE2-QD treated with S-RBD (Fig. 1G, H).

To validate the surface immobilization of ACE2, A549 cells stably expressing ACE2-GFP were treated with S-RBD-QD. Time-lapse recordings of live cells revealed robust and frequent initiation of filopodia, with S-RBD-QD dramatically inducing the rearrangement of the plasma membrane labeled by ACE2-GFP, resulting in membrane elongation at the filopodial tips (Fig. 1I, Movie S2). Genome-wide CRISPR studies in A549 ACE2 cells identified specific actin nucleators, such as Arp2/3 and WASH complexes, as known promoters of rapid branched actin assembly (*19, 20*), most clearly observed in actin comets(*21*). The hypothesized role of these actin regulators is generally linked to endosome entry. Both Arp2/3 and WASH complexes have been previously reported to be at the core of filopodia(*2*). Utilizing S-RBD-QD treatment on A549 ACE2 cells transiently transfected with the actin labeling peptide, the calponin homology domain of utrophin (UtrCH), we can readily capture the colocalization of actin and ACE2 on the filopodia (Fig. 1J, Movie S3).

### Filopodia Mechanosensation Readout by Intracellular Actin Remodeling (or Enzymatic activity of ACE2 drives filopodia and intracellular actin remodeling)

The surface immobilization of S-RBD-QD-ACE2 revealed an unknown role of ACE2 in driving and anchoring filopodia. We first characterized the scale of filopodia induced by this interaction. A custom MATLAB program was developed to integrate both filopodia number and length into one parameter, “Filo-Ratio,” representing the area ratio of filopodia versus other areas of the cell body excluding filopodia at the basal plane (Fig. 2A). Second, since Ang II induced a slowing down of anti-ACE2-QD motility much like S-RBD, we also aimed to test whether the Ang II-ACE2 interaction induced comparable filopodia, as observed with S-RBD. A549 and A549 ACE2 cells were transiently transfected with UtrCH, and Filo-Ratio quantification was obtained following treatment with Ang II or S-RBD. Both treatments induced significant filopodia formation, with a 3.6fold increase in the Filo-Ratio (from 0.05 to 0.18) at the 1-hour time point (Fig. 2B). However, in comparison to Ang II, S-RBD caused faster filopodia formation (5 min) and maintained the persistence of filopodia longer (2 hrs), especially at the basal surface of the cell (Fig. 2C). The filopodia induced by S-RBD exhibit a dose-dependent response (Fig. S3) and are susceptible to drug treatment specifically targeting filopodia formation (Fig. S4).

**Fig. 2.**
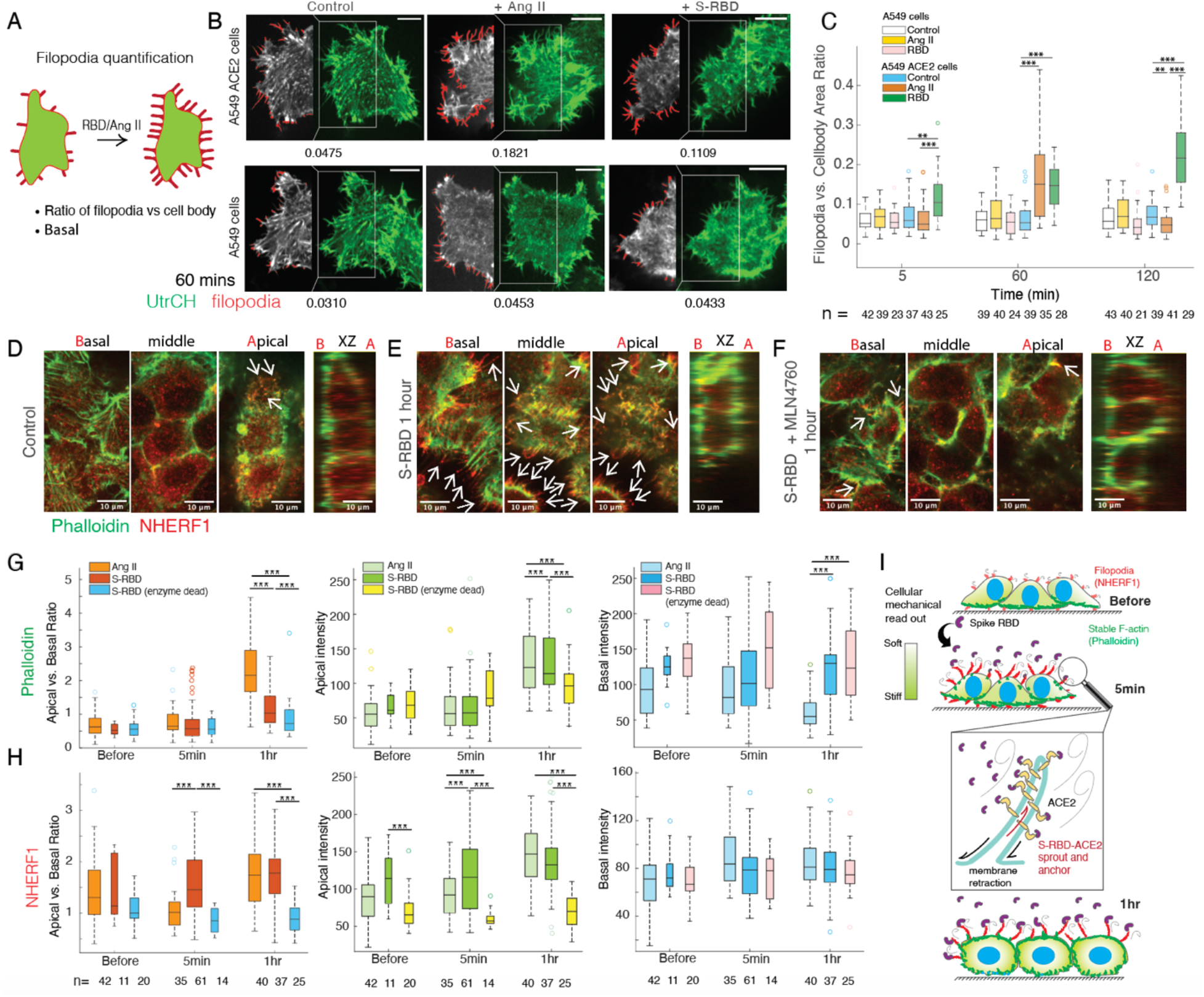
Filopodia Mechanosensation Readout by Intracellular Actin Remodeling. **(A)** Illustration of Filopodia Quantification: Presented as the area ratio of filopodia (red) vs. cell body (green), excluding filopodia, at the basal plane of A549 or A549 ACE2 cells. This allows the extraction of differences in filopodia after treatment with Ang II and S-RBD. **(B)** Representative Images of Filopodia at the Basal Plane: Shown for A549 or A549 ACE2 cells, using transiently transfected UtrCH (green), at the 60-minute time point of treatment from Control, Ang II (200nM), and S-RBD (100nM) treatment. Scale bar is 10 μm. The white box highlights the recognized filopodia area (red) overlaying with the cell (gray). The number of area ratios calculated for each example cell is presented under the image. **(C)** Filopodia vs. Cell Body Area Ratio Plots: Calculated for A549 and A549 ACE2 cells under the conditions of no treatment, Ang II (200 nM), and S-RBD (100 nM) treatment. (**P < 0.01; ***P < 0.001). The number of cells (n) for each calculation is presented under each box. **(D, E, F)** Representative Images of Filopodia Staining: Done by NHERF1 (red) and stable F-actin phalloidin (green) at the basal, middle, apical planes of the x-y axis, and side view of the x-y axis of A549 ACE2 cells, under the conditions of no treatment (D), S-RBD 1 hr (E), and S-RBD + MLN4760 1 hr (F) treatment. White arrows highlight the NHERF1-stained filopodia tips not overlaying with phalloidin staining. Scale bar is 10 μm. **(G, H)** Box Plots of Phalloidin (G) and NHERF1 (H) Intensity: For Basal vs. Apical, apical intensity, and basal intensity calculated from A549 ACE2 cells under the conditions of Ang II (200nM) and S-RBD (100nM) treatment. Additionally, data for A549 ACE2 H374N, H378N (enzyme dead) S-RBD (100nM) treatment cells. Time-lapse of each treatment is calculated for before, 5 minutes, and 1 hour. (**P < 0.01; ***P < 0.001). The number of cells (n) for each calculation is presented under each box. **(I)** Illustration of Intracellular Actin Remodeling: In response to filopodia mechanosensation activated by RBD-ACE2 interaction at the time points of 0 minutes, 5 minutes, and 1 hour of S-RBD 100nM treatment in the A549 ACE2 cells. The green gradient in the cell indicates the degree of the intracellular mechanical readout. The illustration highlights the change in location and sizes of NHERF1-labeled filopodia (red) and phalloidin-stained stable F-actin (green), as well as the degree of the intracellular mechanical readout (green gradient) between 0 to 1 hour duration of S-RBD (purple) treatment.

As previously introduced, filopodia formation implicates the responsiveness of specific cells to signaling cues. It is unclear which external signals ACE2 helps sense by driving filopodia. However, the abundant and persistent filopodia satisfy an unprecedented condition of applying tensile forces through filopodia to the cell, triggering filopodial mechanotransduction. It is crucial to understand downstream cellular responses to this mechanical stimulus, to help identify the upstream sensing signals. We first examined the time-dependent cell adaption by characterizing the actin remodeling in confluent A549 ACE2 cells. We labeled both with the actin marker phalloidin, which prefers stable filamentous actin (F-actin) and stress fibers, we also co-labeled a specific actin probe that binds to ACE2 via the PDZ domain(*22*). This markers also translocates from the cytosol to cortical protrusions when activated(*23*) via the Na+/H+ exchange regulatory factor-1 (NHERF1).

Prior to treatment of Ang II or S-RBD, NHERF1 only overlaps with phalloidin at apparently random apical protrusions(Fig. 2D). After 1 hour of S-RBD treatment, both labeling intensities and the physical overlap of the phalloidin and NHERF1 markers increase significantly. Specifically, NHERF1 labels hair-like longer tips extending out of the phalloidin-labeled fibers throughout the cell (Fig. 2E). The side view of the treated cells reveals a pronounced increase in the F-actin signals on the apical side of the cell, where Ang II or S-RBD is administered. Treatment with the ACE2 inhibitor MLN4760 together with S-RBD abolishes these changes (Fig. 2F). To quantify the actin remodeling, we measured the phalloidin and NHERF1 intensities on the apical and basal sides of cells from the side view of z-stack confocal images obtained from Ang II or S-RBD treated A549 ACE2 cells, and the S-RBD treated A549 H374N, H378N ACE2 cells (enzymatic dead mutations). For each condition, the measurements are recorded for before, the fast (5 min), and slow (1 hour) time points.

For confluent cells, the overall F-actin signal increase was most pronounced at the apical side of the cells (Fig. 2G). NHERF1 showed a fast, persistent, and ∼two-fold increase in intensity following S-RBD treatment (Fig. 2H). Phalloidin intensities matched the increase of NHERF1 at a slower pace by the 1-hour time point. Ang II induced both NHERF1 and Phalloidin intensity changes comparable to those of the S-RBD, but they also at a slower rate.

To test the importance of ACE2 activity for rearrangement of actin, we compare cells expressing wild type ACE2 to cells expressing a catalytically dead version of ACE2 with H374N, H378N mutations in the active sites. No significant changes were measured for either NHERF1 or phalloidin intensities were measured in S-RBD treated A549 H374N, H378N ACE2 cells compared to wild type ACE2. Combining the results of activator Ang II and enzyme dead mutation conditions of ACE2, we confirmed that both filopodia-driven and intracellular remodeling capabilities of ACE2 depend on its enzymatic activities.

In summary, our observations indicate that the introduction of Ang II or the S-RBD elicits a rearrangement of cellular activity, leading to the development of a denser cortex at the specific loci where ACE2 drives and accumulates at the filopodia. This cellular response suggests an analogy to the cellular interpretation of the administered Ang II/S-RBD environment, potentially akin to the cellular response to increased mechanical loads (Fig. 2I).

## Filopodia mechanosensation readout by cell bulk stiffness

To test our hypothesis concerning the cellular response to mechanical deformation of filopodia, wherein a thicker actin cortex assembles to resist deformation, we proceeded with the assessment of cell bulk stiffness utilizing microrheology based on multi-particle tracking. The microrheology method facilitates the determination of elastic and viscous moduli by analyzing the trajectories of probes, such as beads or endosomes, within the *in vitro* actin polymer solution or cytosol(*24*). To acquire trajectories of the QD endosomes, we focused on the endocytosed S-RBD-QD at the centrosomal plane, distinct from the surface immobile portion located at the basal plane (Fig. 3A).

**Fig. 3.**
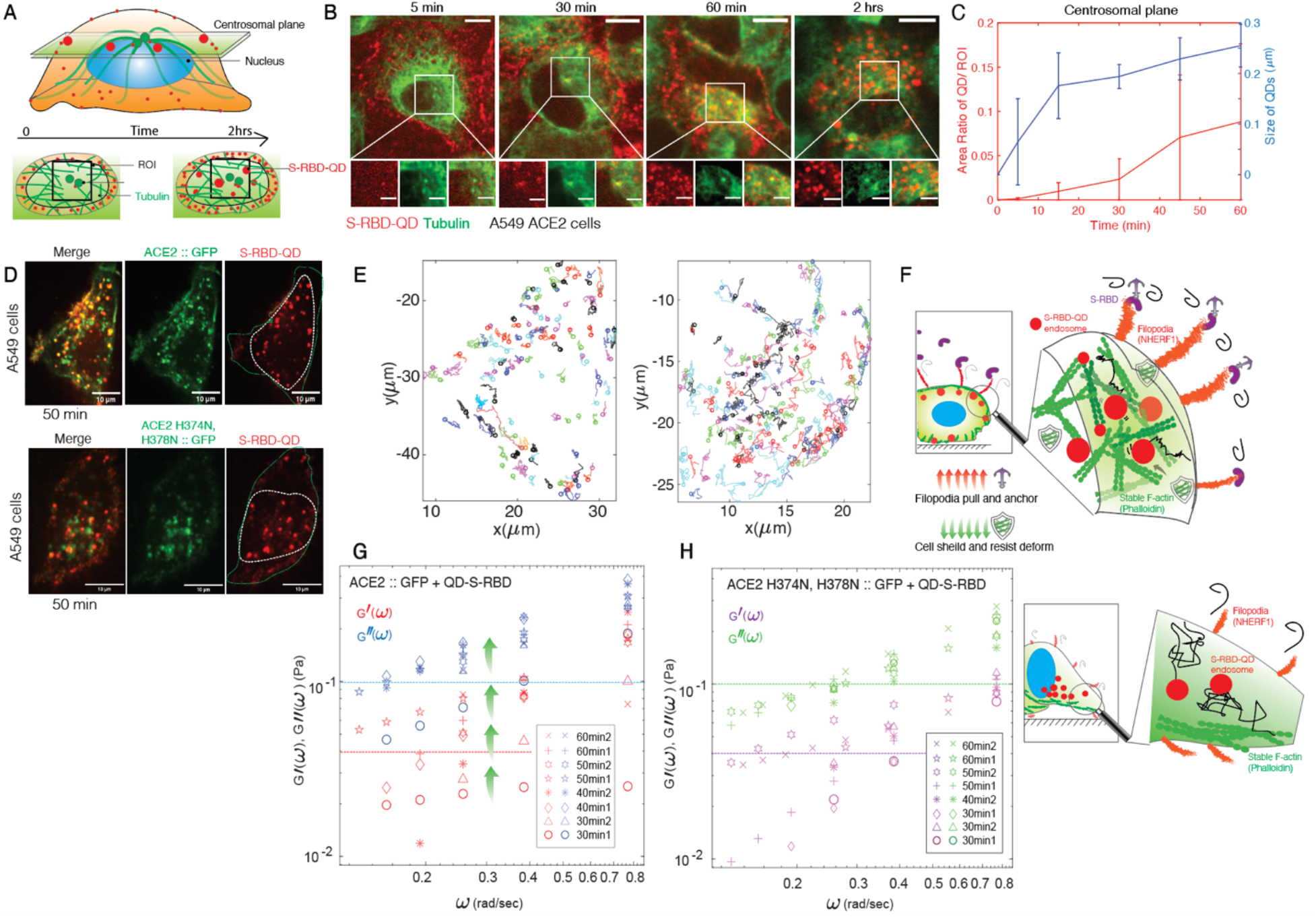
Filopodia Mechanosensation Readout by Cell Bulk Stiffness. **(A)** Illustration of Capturing Time-lapse of S-RBD-QD Endosomes: During 0 to 2 hours at the centrosomal plane of A549 ACE2 cells. A square with a side length of 15 μm centered at the centrosomes is selected as the region of interest. **(B)** Representative Images of S-RBD-QD Endosomes (Red): Shown in the whole cell view (Scale bar is 10 μm) and at the enlarged ROI selected (Scale bar is 5 μm) after 5, 30, 60, 120 minutes of treatment. Tubulin (green) staining shows the location of the centrosomes. **(C)** Area Ratio of QD/ROI (Red) and Size of QDs (Blue) Plots: In a time-lapse of 5, 15, 30, 45, 60 minutes after S-RBD-QD treatment (S-RBD 100nM) on A549 ACE2 cells from the ROI at the centrosomal plane. **(D)** Representative Images of S-RBD-QD Endosomes: In A549::ACE2 GFP/ A549::ACE2 H374N H378N GFP cells after treatment of S-RBD-QD for 50 mins. In the red channel, the green dash line shows the cell boundary lined by ACE2-GFP, and the white dash line shows the QD tracking region selected for trajectory calculation. Scale bar is 10 μm. **(E)** Tracking Trajectories: In the duration of 2 minutes of S-RBD-QD endosomes as captured in the ROI shown in (D). **(F)** Illustration of Cell Bulk Stiffness Increase on A549::ACE2 GFP/ A549::ACE2 H374N H378N GFP cells: Driven by filopodia formation induced by S-RBD-ACE2 interaction. The persistent pulling of filopodia (NHERF1) anchored by S-RBD at the tip causes mechanical responses in the cell bulk in the form of stable F-actin (phalloidin) shielding to resist deformation. The increased resistance caused by F-actin polymerization hinders endosome diffusion in the cytosol, which can be used to extract viscoelastic medium properties. **(G)** Plots of Elastic (G’, Red) and Viscous (G’’, Blue) Modulus versus Frequency (ω): Calculated from S-RBD-QD endosomes in ACE2-GFP stable expression A549 cells after 30 min (circles, triangles), 40 min (diamonds, stars), 50 min (pentagons, hexagons), and 60 min (pluses and crosses) of treatment. Green arrows highlight the trend of continuous increase in the elastic modulus over time. **(H)** Plots of Elastic (G’, Purple) and Viscous (G’’, Green) Modulus versus Frequency (ω): Calculated from S-RBD-QD endosomes in ACE2 H374N H378N-GFP stable expression A549 cells after 30 min (circles, triangles, diamonds), 40 min (stars), 50 min (pluses and hexagons), and 60 min (pentagons and crosses) of treatment.

From a square region centered at the centrosomes with a side length of 10 μm, we carefully examined the speed of endocytosis induced by the interaction between S-RBD-QD and ACE2 (Fig. 3B). Notably, the sizes of the QD trajectories recorded at the centrosome plane exhibited a sudden increase approximately 15 minutes post the S-RBD-QD treatment, stabilizing thereafter at a mean radius of approximately 0.25 μm. This behavior contrasts with a more gradual increase observed in QD sizes at the basal plane (Fig. S2). Importantly, this distinctive change in the size of the QD-defined domain remains consistent across various treatments, including S-RBD-QD dosage variations, drug interventions targeting filopodia, and modulation of ACE2’s enzymatic function (Fig. S5). Furthermore, an analysis of the endosome population, as indicated by the area of the QD in relation to the Region of Interest (ROI), reveals a phase of gradual increase preceding the 30-minute mark after S-RBD-QD treatment, followed by a more rapid increase thereafter (Fig. 3C). Both the speed and final count of QD-marked endosomes exhibit a correlation with the dosage of S-RBD-QD treatment, filopodia formation, and the enzymatic activity of ACE2 (Fig. S5).

To investigate changes in the mechanical properties of the cytosol, we systematically recorded time-lapse videos of S-RBD-QD endosomes at 10-minute intervals, spanning the 30 to 60-minute post-treatment period on A549 cells expressing either ACE2-GFP or ACE2 H374N, H378N – GFP (Fig. 3D). To precisely capture a representative population of newly endocytosed quantum dots (QD), we excluded immobile ACE2 accumulations on the cell surface by confining particle tracking to a selected region encompassing the cytosol.

The trajectories of endocytosed QD endosomes at various time points following treatment revealed constrained motion within wild type ACE2-GFP-containing cells, contrasting with the behavior observed in cells with mutant catalytically dead ACE2 (Fig. 3E, Movie 4,5). Furthermore, trajectories were predominantly situated at the periphery of ACE2-GFP cells, as opposed to a more even distribution throughout the entire region observed in cells with ACE2 mutations (Fig. 3E). Notably, the absence of ensemble retrograde centrifugal motions for QD endosomes in ACE2-GFP cells eliminates the possibility of active transport (Movie 4, 5; overlay trajectories showing retrograde or antegrade displacement from centrosomes). The mean square displacement demonstrated linearity (data not shown), suggesting that the observed confinement is not dynamic and not permanent. Building upon these observations, our working hypothesis is that ACE2 exerts an influence on the mechanical properties of the cytosol by orchestrating remodeling of intracellular actin. This, in turn, results in a stochastic and extensive hindrance on QD endosomes (Fig. 3F). From the trajectories obtained, we conducted a comprehensive analysis to derive the elastic (G’) and viscous (G’’) moduli in relation to frequency (ω) through one-particle microrheology. Notably, both moduli exhibited a conspicuous elevation in ACE2-GFP cells. A surprising revelation emerged as the elastic modulus in ACE2-GFP cells displayed a time-dependent increase, indicating an escalating demand for energy input to induce deformation. On the contrary, the viscous modulus in ACE2-GFP cells remained relatively consistent throughout the observation period.

The observed stiffening of the cellular bulk in ACE2-GFP cells is robustly correlated with filopodia anchoring mediated by ACE2. This inference is substantiated by the observation that the measured time courses of both elastic (G’) and viscous (G’’) moduli in ACE2 mutation cells remained consistent, exhibiting no significant changes. The observation of alterations in bulk cell stiffness serves as a pivotal insight that supports the proposed role of ACE2 in mediating the cellular perception of environmental mechanical loads.

## Concurrent intracellular calcium rise

Mechanosensitive ion channels, exemplified by Piezo1 and TRPV4, have been identified to localize on filopodia and exhibit responsiveness to mechanical stimuli(*25*). Prolonged intracellular calcium elevation, concomitant with the upregulation of Piezo1 expression, has been observed following treatment with S-RBD in lung vascular endothelial cells (*15*). In our investigation of calcium signaling in ACE2-mediated filopodia mechanosensation, we transiently transfected the live-cell calcium indicator Gcamp6f and measured cytosolic Gcamp6f intensity in A549 and A659 ACE2 cells under the treatment of Ang II and S-RBD at 5 minutes and 1-hour time points.

Significantly, only in ACE2-expressing cells, a robust intracellular calcium rise was detected at 5 minutes for S-RBD, with comparable calcium levels persisting at 1 hour. In contrast, Ang II induced a slower and lower increase in calcium levels compared to S-RBD (Fig. 4A, B). The dose-dependence of the calcium increase induced by both Ang II and S-RBD was confirmed (Fig. S6). Moreover, the ACE2 inhibitor DX600 effectively eliminated the increase in intracellular calcium induced by both Ang II and S-RBD (Fig. 4C).

**Fig. 4.**
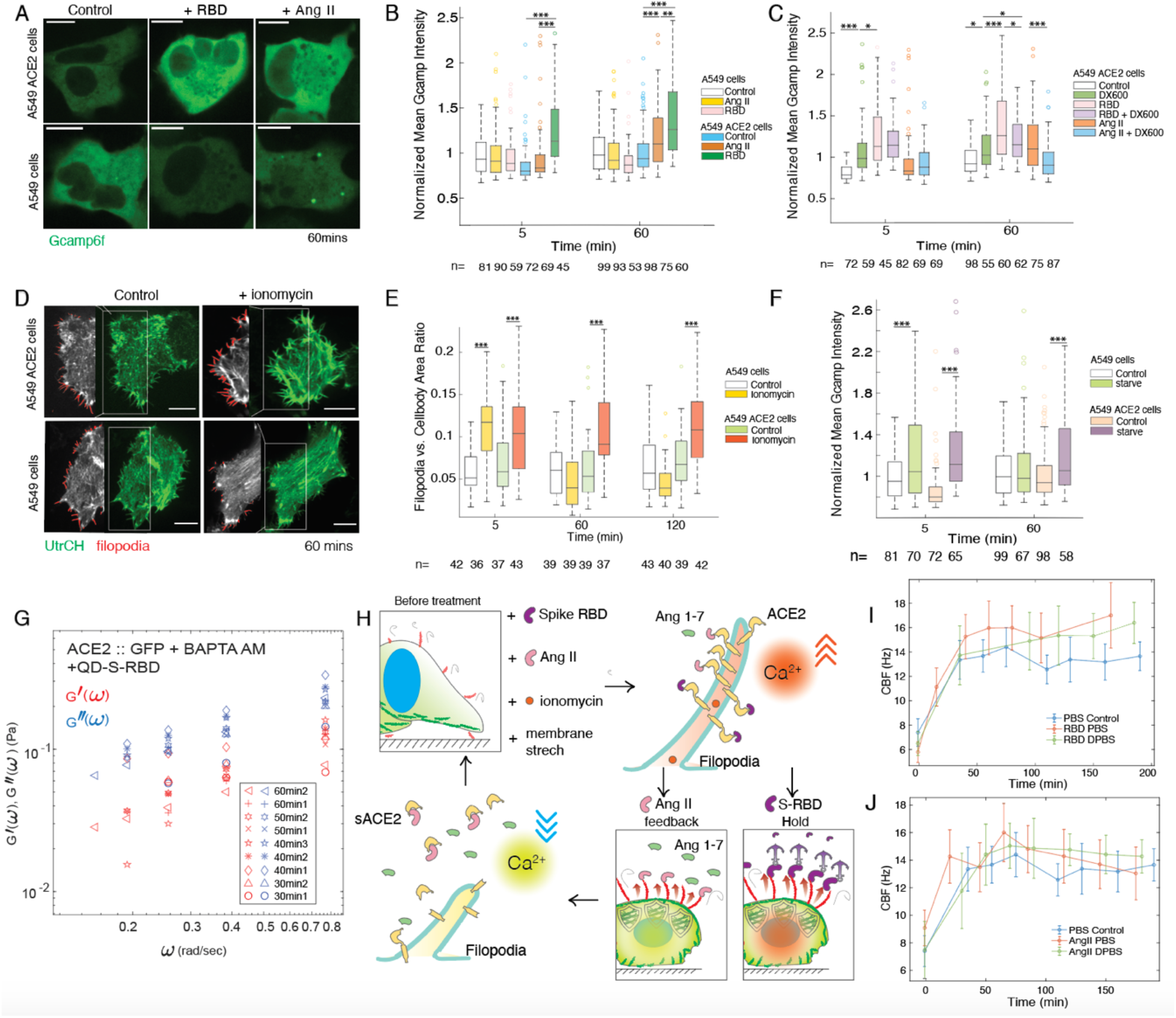
Concurrent Intracellular Calcium Rise and ACE2 mediated filopodial mechanosensation model. **A)** Representative Images of Intracellular Calcium Levels: Live A549 and A549 ACE2 cells were transiently transfected with Gcamp6f and imaged at the 60-minute time point after Ang II and S-RBD treatment. **(B, C)** Box Plots of Normalized Mean Gcamp6f Intensity: Time-lapse of each treatment was calculated for 5 and 60 minutes (*P < 0.1; ***P < 0.001). Each measured intensity is normalized by the mean intensity of A549 ACE2 cells under control conditions. The number of cells (n) for each calculation is presented under each box. (B) Measured intensity of the cell body from A549 and A549 ACE2 cells under the conditions of control, Ang II (200nM), and S-RBD (100nM) treatment. (C) Measured intensity of the cell body from A549 ACE2 cells under the conditions of control, DX600 (100nM), S-RBD (100nM), DX600 + S-RBD, Ang II (200nM), and DX600 + Ang II treatment. **(D)** Representative Images of Filopodia at the Basal Plane: Shown for A549 or A549 ACE2 cells, using transiently transfected UtrCH (green), at the 60-minute time point of treatment from Control and ionomycin (0.25μM) treatment. Scale bar is 10 μm. The white box highlights the recognized filopodia area (red) overlaying with the cell (gray). **(E)** Filopodia vs. Cell Body Area Ratio Plots: Calculated for A549 and A549 ACE2 cells under the conditions of no treatment and ionomycin (0.25 μM) treatment (***P < 0.001). The number of cells (n) for each calculation is presented under each box. **(F)** Box Plots of Normalized Mean Gcamp6f Intensity: Time-lapse of glucose+serum starvation treatment was calculated for 5, and 60 minutes (***P < 0.001). Each measured intensity is normalized by the mean intensity of A549 cells under control conditions. The number of cells (n) for each calculation is presented under each box. **(G)** Plots of Elastic (G’, Red) and Viscous (G’’, Blue) Modulus versus Frequency (ω): Calculated from S-RBD-QD endosomes in ACE2-GFP stable expression A549 cells after 30 min (circles, triangles), 40 min (diamonds, stars, pentagons), 50 min (crosses and hexagons), and 60 min (pluses and inverted triangles) of treatment with S-RBD-QD and BAPTA AM. **(H)** Illustration of the Model of Filopodia Mechanosensation Mediated by ACE2: Filopodia spout and anchor in response to calcium rise in the cytosol triggered by S-RBD/Ang II/ionomycin/or other factors that result in membrane stretch. The adherent cells reorganize their F-actin and polymerize F-actin at the cortex as shields from deformation. Intracellular calcium rises to a threshold to induce ACE2 hydrolysis and detach its catalytic pocket from the plasma membrane. S-RBD holds the filopodia; while the Ang 1-7 produced on the membrane or externally by soluble ACE2 decreases intracellular calcium rise and filopodia, returning the cell to a before-activated status. **(I)** Plot of CBF for HNE Cells: Mean CBF of the ROI selected (details please see the Methods) measured under the condition of adding 50 μl PBS control, RBD 0.3ug in PBS, and RBD 0.3ug in DPBS. Time-lapse of each treatment is calculated for time points between 0 to 3 hrs. **(J)** Plot of CBF for HNE Cells: Mean CBF of the ROI selected (details please see the Methods) measured under the condition of adding 100 μl PBS control, Ang II 200nM in PBS, and RBD 1μg in DPBS. Time-lapse of each treatment is calculated for time points between 0 to 3 hrs.

The measured response from intracellular calcium elevation aligns well with filopodia growth. We next explored whether the two effectors synergize with each other. First, we tested the filopodia response induced by ionomycin. In A549 cells, filopodia increased significantly and rapidly, but the increase did not persist until the 1-hour time point. In contrast, in A549 ACE2 cells, the filopodia induced by ionomycin remained at a relatively high level (Filo-Ratio ∼0.1 compared to ∼0.18 induced by S-RBD) in a time window from 5 minutes to 2 hours (Fig. 4D, E, Fig. S7A, B). Secondly, we used glucose and serum starvation to induce filopodia. We measured a significant increase in intracellular calcium rise in both A549 and A549 ACE2 cells in response to the starvation treatment at the 5-minute time point. We also observed a similar trend of prolonged high calcium levels in A549 ACE2 cells and a drop in calcium levels in A549 cells at the 1-hour time point (Fig. 4F).

In conclusion, we confirmed that ACE2-induced massive and persistent filopodia could bypass the enzymatic activators and be directly triggered by intracellular calcium rise. We further validated the role of calcium in filopodia mechanosensation through time-lapse measurement of cell bulk stiffness under the treatment of S-RBD and BAPTA AM. We confirmed that the mechanical change in the cytosol caused by S-RBD is abolished by quenching the calcium rise (Fig. 4G, Fig. S7C).

## Discussion

ACE2 exhibits a ubiquitous expression across various human tissues, including the small intestine, kidney, heart, adipose tissue, and lungs(*26*). Beyond its well-established roles in blood pressure regulation within the renin–angiotensin system (RAS) pathway and neutral amino acid transport in the intestine, the broader physiological functions of ACE2 in these diverse tissues including the airway epithelium and remain less clear. The evolution of specific Corona viruses like SARS-CoV-2 to use ACE2 as its receptor hardly appears to be an accident and yet we do not specifically understand how the interactions with viral interactions with ACE2 expressing cells influence either the vasculature or other key tissues. Despite our earlier observations of ACE2 in airway epithelia, the role of ACE2 in airways remains unclear. A meticulous examination of the detailed distribution of the ACE2 protein in these tissues is conducted to evaluate susceptibility to SARS-CoV-2 virus entry. A notable characteristic is the prominent and significant staining of ACE2 along the inner surfaces of liquid-contacting luminal structures that undergo contractile-relax motions(*27*). This includes structures in enterocytes, kidney tubules, gallbladder, pancreas duct, nasal or bronchus epithelia, as well as vascular endothelia. In this study, we unveil a novel and pertinent role of ACE2 as a mediator of filopodia mechanosensation, wherein its activation elicits multiple cellular responses, including actin remodeling, stiffening of cytosol mechanics, and a simultaneous intracellular increase.

Our discovery of ACE2’s role was illuminated through a comparative analysis of the signaling pathways triggered by one of ACE2’s best-known enzymatic substrates, Ang II, and the S-RBD. We systematically assessed the impact of these ligands on ACE2’s mobility, induced filopodia, and intracellular calcium rise. While both Ang II and S-RBD induced filopodia and intracellular rise, contingent on the catalytic activities of ACE2, notable distinctions emerged. Ang II led to filopodial retraction at the two-hour time point and elicited a lower calcium rise at a dose equivalent to that triggering comparable filopodial scales as S-RBD. In contrast, S-RBD exhibited no filopodial inactivation at the two-hour time point, inducing both swifter and more prolonged intracellular rise. Furthermore, our investigation revealed that prolonged ACE2 activation prompted a cellular response with a mechanical component to resist deformation, potentially influencing its intracellular trafficking within the cytosol. Consequently, this sustained state may not be conducive to cellular health if perpetuated indefinitely.

In our search for a potential exit mechanism from how ACE2 rearranges following Ang II stimulation, we identified two candidate pathways: shedding of the ACE2 catalytic ectodomain (referred to as soluble ACE2 or sACE2) triggered by intracellular calcium rise, and the potential influence of the vasodilator angiotensin 1-7 (Ang 1-7) generated by the Ang II-ACE2 interaction. While the link between cytosolic calcium and sACE2 shedding is well-documented, we conducted experiments to test the treatment of Ang 1-7 together with S-RBD on A549 ACE2 cells. Our findings confirmed that the treatment of Ang 1-7 together with S-RBD led to a decrease in intracellular calcium to levels lower than before treatment. Moreover, S-RBD did not induce filopodia formation when treated together with Ang 1-7 (Fig. S7D, E, F). The revelation of Ang 1-7 as a possible mechanism for downregulation, especially as a downstream product from ACE2 enzymatic activity, informs our hypothesis of a sustainable model for filopodia mechanosensation orchestrated by ACE2 (Fig. 4H).

External stimuli, such as enzymatic substrates of ACE2 or membrane stretch opening mechanosensitive ion channels, initiate filopodia formation with ACE2 accumulating and becoming immobile on them. The massive and persistent growth of filopodia, facilitated by ACE2 dragging the plasma membrane, conveys a message of increased load to the cells, prompting them to respond by reinforcing the cytoskeletal defense and resisting deformation. The pulling of the filopodial membrane also leads to an elevation in intracellular calcium levels, reaching a threshold that activates sACE2 shedding. As sACE2 shedding occurs, the enzymatic ectodomain of ACE2 is detached, diminishing the power to sustain filopodia. In cases where ACE2 is activated by Ang II, the increase in Ang 1-7, newly generated by ACE2 on the membrane or nearby shredded sACE2, accelerates filopodia retraction. Consequently, the cell completes a cycle for response to environmental cues.

To rigorously test this hypothesis, we turned to the air-liquid-interface (ALI) cultured human airway epithelial cells, which we previously revealed to exhibit pronounced microvilli phenotypes in response to SARS-CoV-2 virus infection(*1*). In this study, we treated the cells with S-RBD and Ang II and recorded their cilia beating frequency (CBF) as a readout for the cells’ response to external mechanical loads. We observed an increase in CBF for both S-RBD (Movie S6) and Ang II treatments, with the amplitude of CBF change dependent on environmental calcium concentrations (Fig. 4 I, J, Fig. S8 E-I). Of note, Ang II exhibited a return of CBF to control levels at the 3-hour time point, while S-RBD maintained a prolonged high level of CBF at the same time point.

In conclusion, our findings support the existence of an Ang II-ACE2-filopodia-calcium-Ang 1-7 loop that may also apply to the mechanical response of airway epithelial cells. It is important to note that there is currently no reported existence of Ang II in the mucus. Further research is needed to confirm the constitution of Ang II in mucus or to identify other possible airway-specific enzymatic substrates for ACE2, which functions similarly to Ang II. Biochemical studies, aiming to identify substrates for recombinant soluble human ACE2, have revealed that ACE2 hydrolyzes various substrates, including angiotensin I, 1-9, II, 1-7, 1-5; apelin-13, 36; bradykinin, Neurotensin 1–8, etc(*28*). Our study not only encourages the characterization of whether other substrates induce a similar response of filopodia mechanosensation through ACE2 catalytic activities but also suggests the potential discovery of novel mechanical regulations relevant to the local environment in diverse tissues where these substrates are uniquely distributed.

These enzymatic substrates of ACE2 hold the potential to facilitate the development of novel signal peptides aimed at driving tissue-specific cell migration in wound healing or stem cell differentiation. Understanding the function of ACE2 in host cells will also provide valuable information for COVID treatment strategies, particularly in addressing mechano-pathologies related to the dysregulation of cells’ environmental sensations, such as fibrosis or apoptosis. Therapeutic targets inspired by the signaling cascade’s exit mechanism hold significant promise for treatment development.

## Acknowledgments

We thank Dr. T. Kanie for sharing the lentivirus compatible vector and protocol for generating stable cell lines. We thank Dr. Ran Cheng for helpful discussions.

## Author contributions

Conceptualization: WH, CTW, PKJ; Methodology: WH, CTW; Investigation: WH, CTW; Visualization: WH; Funding acquisition: PKJ; Project administration: PKJ; Supervision: PKJ; Writing – original draft: WH; Writing – review & editing: WH, CTW, PKJ

## Data and materials availability

The datasets to be analyzed during the current study will be available from the corresponding author on reasonable request.

## Supplementary Materials

Materials and Methods

Figs. S1 to S8

Movies S1 to S6

