## Supplementary material for "Filopodial Mechanotransduction is regulated by Angiotensin-Converting Enzyme 2 (ACE2) and by SARS-CoV-2 spike protein": Materials and Methods Figs. S1 to S8

**The PDF file includes:**

Materials and Methods
Figs. S1 to S8

**Other Supplementary Materials for this manuscript include the following:**

Movies S1 to S6

### Materials and Methods

#### Plasmids

The Gateway entry vector for Homo sapiens ACE2 without a stop codon was purchased from Dharmacon (Entrez Gene 59272). Quick-Change mutagenesis was applied to generate ACE2 H374N and H378N mutations. Complementary forward and reverse primers with a point mutation at the mutation site in the middle were used for PCR. The PCR product was treated with 20U of DpnI at 37°C for 1 hour and then transformed into competent cells.

The Gateway cloning-compatible lentiviral vector, pWPXLd/LAP-C/puro/DEST vector, was previously described (cite T. Kanie et al., 2017). The pWPXLd vector was a gift from Prof. Didier Trono (Addgene plasmid #12258). The pWPXLd/LAP-C/blast/long EF/DEST vector was created by inserting the DEST/C-terminally LAP tag/blasticidin resistance cassette into the pWPXLd vector. The lentivirus envelope and packaging vectors, pCMV-VSV-G and pCMV-dR8.2 dvpr, were gifts from Prof. Bob Weinberg (Addgene plasmid #8454 and #8455). Lentiviral vectors containing ACE2 or ACE2 H374N and H378N mutations were generated by LR recombination between ACE2 entry vectors without a stop codon and the pWPXLd/LAP-C/blast/long EF/DEST vector.

#### Cell culture, drug treatment, and transfection

The A549 cells were cultured in DMEM medium (Gibco, 12800017) containing 10% FBS (Gemini, 100-106), 100 U/mL Penicillin-Streptomycin (Thermo Fisher Scientific, 15140163) at 37°C in 5% CO2. The ACE2 or ACE2 H374N, H378N stable cell lines were generated using Lentivirus. 500 ng of Lentivirus vectors containing the ACE2 or ACE2 H374N, H378N gene were transfected together with 150 ng of pCMV-VSV-G, 350 ng of pCMV-dR8.2 dvpr using the transfection reagent, 3 µl of Fugene 6 (Promega, E2692). After 48 hours, the virus was filtered with a 0.45 µm PVDF filter (Millipore, SLHV013SL) and mixed with a 4-fold volume of fresh media containing 12.5 µg/ml polybrene (Millipore, TR-1003-G). After incubating the virus with A549 wt cells for 66 hours, 10 µg/ml blasticidin (Corning, 30-100-RB) was applied with fresh culture medium to select the transfected cells for 5 days.

Drug treatment was performed with 100nM, 200nM, 400nM Angiotensin II (Sigma-Aldrich, A9525); 75nM, 100nM, 150nM Spike-RBD (BPS Bioscience, 100937-2); 1nM, 3nM MLN4760 (Sigma-Aldrich, 5.30616); 10µM DX600 (Cayman, 22186); starvation medium DMEM, no glucose (Thermo Fisher Scientific, 11966025); 3 µM LY294002 (Med Chem Express, HY-10108); 0.25 µM ionomycin (MP Biomedicals, No. 15507001); 5 mM BAPTA AM (Life Technologies, B-6769); 200nM Angiotensin 1-7 (apexbio, A1041); 2 µg/ml soluble ACE2 (Boster, RCOV09).

For transient transfection, cells were seeded in glass-bottom chambers for 24 hours and then transfected with the respective DNA plasmids using Lipofectamine™ 2000 Transfection Reagent (Invitrogen, 11668-027). The plasmids used in this work are pGP-CMV-GCaMP6f, Utr-CH-EGFP (Addgene, No. 26737).

#### Primary human nasal cell culture

Epithelial cultures were generated using an already well-established protocol in our laboratories (Vladar et al., 2016). After obtaining informed consent (Stanford IRB protocol #42710), subjects underwent brushing of the inferior turbinate from both nasal cavities to obtain a cell sample. The sample was immediately transferred to the laboratory and dissociated in a 1:1 mixture of proteolytic enzymes (Accutase, Thermo Fisher Scientific Inc.) and nonenzymatic cell dissociation solution (C5914, Millipore-Sigma Inc.) for 10 minutes at 37ºC. After gentle agitation the cells were pelleted, resuspended in basal cell proliferation media (PneumaCult-Ex Plus medium, STEMCELL Technologies, Vancouver, Canada) supplemented with antibiotics and plated on collagen-coated Transwell inserts (0.33 cm2) at a density of 20,000 cells per insert with media added to both the basal and apical sides. Once the cells were fully confluent, the media was replaced with PneumaCult-ALI medium (STEMCELL Technologies, Vancouver, Canada) in the basal chamber and the apical surface exposed to provide an air liquid interface (ALI). Monolayers were grown at ALI for an additional 3 weeks to promote differentiation into a nasal epithelium with basal, multiciliated and secretory cells.

Live-cell labeling of QD to ACE2

For adherent A549 cells cultured for imaging, the cells were seeded on 4-well glass-bottom chambers in 35mm dishes(Greiner Bio-One, No. 627870) coated with fibronectin (EMD Millipore, No. FC010).

To label the ACE2 with anti-ACE2-QD in one 35 mm dish, 2 μg of Rabbit Anti-ACE2 antibody (ProSci, 3227) is applied to 400 μl of culture medium and incubated with the cells for 30 mins in the incubator. After the primary antibody is washed three times with culture medium at room temperature, 1 μl of F(ab')2-Goat anti-Rabbit IgG (H+L) Secondary Antibody, Qdot 585 (Thermo Scientific, Q-11411MP), is applied to the cells, together with 1 μl of 50nM SiR-tubulin (Cytoskeleton Inc., CY-SC002) in 400 μl of culture medium. The mixture of QD and SiR-tubulin is incubated with cells for 60 mins and then taken for imaging. Ang II or S-RBD is added after 60 min of incubation if necessary.

To target ACE2 with S-RBD-QD in one 35mm dish, Biotin-tagged RBD (BPS Bioscience, 100937-2) was dissolved in 200 μl of culture medium at a concentration of 200nM and incubated with 1 μl of QD (Thermo Scientific, Q10113MP, 1 μM) for 15 mins. 1 μl of 50nM SiR-tubulin is added to the mixture solution before treatment to cells. For each well, 50 μl of the mixture solution is added to 50 μl of culture medium to a final volume of 100 μl. The dish is immediately taken for imaging. Drugs are added together with the QD mixture solution if necessary.

Immunofluorescence

To detect whole-cell actin remodeling, A549 ACE2 cells, A549 ACE2-GFP cells, and A549 H374N, H378N-GFP cells were grown on 12-mm round coverslips and fixed with 4% paraformaldehyde (AlfaAesar 433689M) diluted in PBS for 15 minutes. The cells were then permeabilized with 0.1% Triton X-100 (Fisher Scientific, AC21568-2500) for 10 minutes at room temperature. Samples were blocked with 5% normal donkey serum (Jackson ImmunoResearch, 017-000-121) in PBS for 1 hour at room temperature. Subsequently, samples were incubated with the primary antibody in PBS at 4 degrees overnight, followed by five washes with PBS. Next, samples were incubated with a fluorescent-labeled secondary antibody for 1 hour at room temperature, followed by five washes with PBS. Finally, coverslips were mounted with SlowFade Antifade Reagent (Life Technologies, S36936) onto glass slides for image acquisition. The antibodies used were as follows: Anti-NHERF-1 (A-7, Santa Cruz Biotechnology, sc-271552), Alexa Fluor™ 488 Phalloidin (Fisher Scientific, No. A12379).

Confocal imaging

For live-cell imaging of QD-labeled ACE2 in A549 cells or recording cilia beating frequency with HNE cells, we utilized a scanning disk confocal microscope (Zeiss Observer Z1, Zeiss; Laser Stack, 3i; CSU-22 confocal scanner unit, Yokogawa) equipped with a Zeiss planapochromat 63x 1.4 oil objective on a vibration-dampened table and a Prime 95B Scientific CMOS camera (Teledyne Photometrics). Images and videos were acquired at a resolution of 0.169 um per pixel in an area of 536 X 544 pixels using Slidebook 6 software (3i). Fluorescent videos were obtained with an exposure time of 100 ms for each channel at a frame rate of 0.9 ~ 1.3 Hz for 60-120 frames. To obtain QD probed ACE2 on the cell surface, the focus plane is set at the basal plane (Fig. 1E, S2B). To obtain QD-ACE2 endosomes in the cytosol, the focus plane is set at the centrosomal plane as indicated in the tubulin staining (Fig. 3A).

To obtain the cilia beating frequency, the transwell insert with HNE cells was transferred to the inner wells of the glass-bottom 24-well dish (GREINER BIO-ONE, 662892) with 500 µl PneumaCult-ALI medium in the well. Ang II, S-RBD, or EP4 agonist was applied with 100 µl PBS/calcium-free PBS to the insert, directly contacting the apical surface of HNE cells (Fig. S8 E, F). Bright-field videos of HNE cells were obtained with an exposure time of 10 ms at a frame rate of 64 Hz. All imaging was conducted at 37°C and 5% CO2 in a live-supporting chamber mounted on the microscope stage.

To acquire images of the whole-cell actin remodeling in fixed samples, a z-stack image was obtained using the same confocal settings as described above, covering the apical and basal range in a step size of 0.5 µm.

<u>Single particle tracking and Microrheology measurement.</u>

The single-particle tracking method, based on a custom MATLAB program generated previously to track transmembrane proteins (citation), is applied to QD-labeled ACE2. In simple terms, the raw image is initially denoised by removing the uneven illuminated background. Next, precise locations of each QD particle are calculated using a band-pass filter. Since both the surface and endosomes of QD show time-dependent augmentation features, we made custom adjustments to the QD recognition section of the algorithm, broadening the range of the size of QDs from single QD of < 0.05 µM to QD aggregations of < 0.25 µM. Following this, a region of interest (ROI) mask excluding the immobile QD portions merged on the cell surface is created by hand drawing. The trajectories in the ROI are calculated using the minimum distance algorithm (citation) and checked by overlaying the trajectories with the original videos (Movie S4, S5). The elastic and viscous modules are calculated using a previously described one-particle microrheology algorithm (citation) using GDL (GRAPHISOFT).

<u>Cilia Beating Frequency CBF</u>

To calculate the CBF from the bright-field videos of live HNE cells on the confocal microscope, a previously reported method based on MATLAB (citation needed) is slightly adjusted to enable customized region-of-interest selection by hand drawing. The adjustment is made to confine the measurement of CBF to individual ciliated HNE cells, decrease the batch-to-batch heterogeneity of the ciliated cell density and thus greatly increasing the precision of the CBF measurement by excluding background cells that are not ciliated (Fig. S8, A, B, C, D).

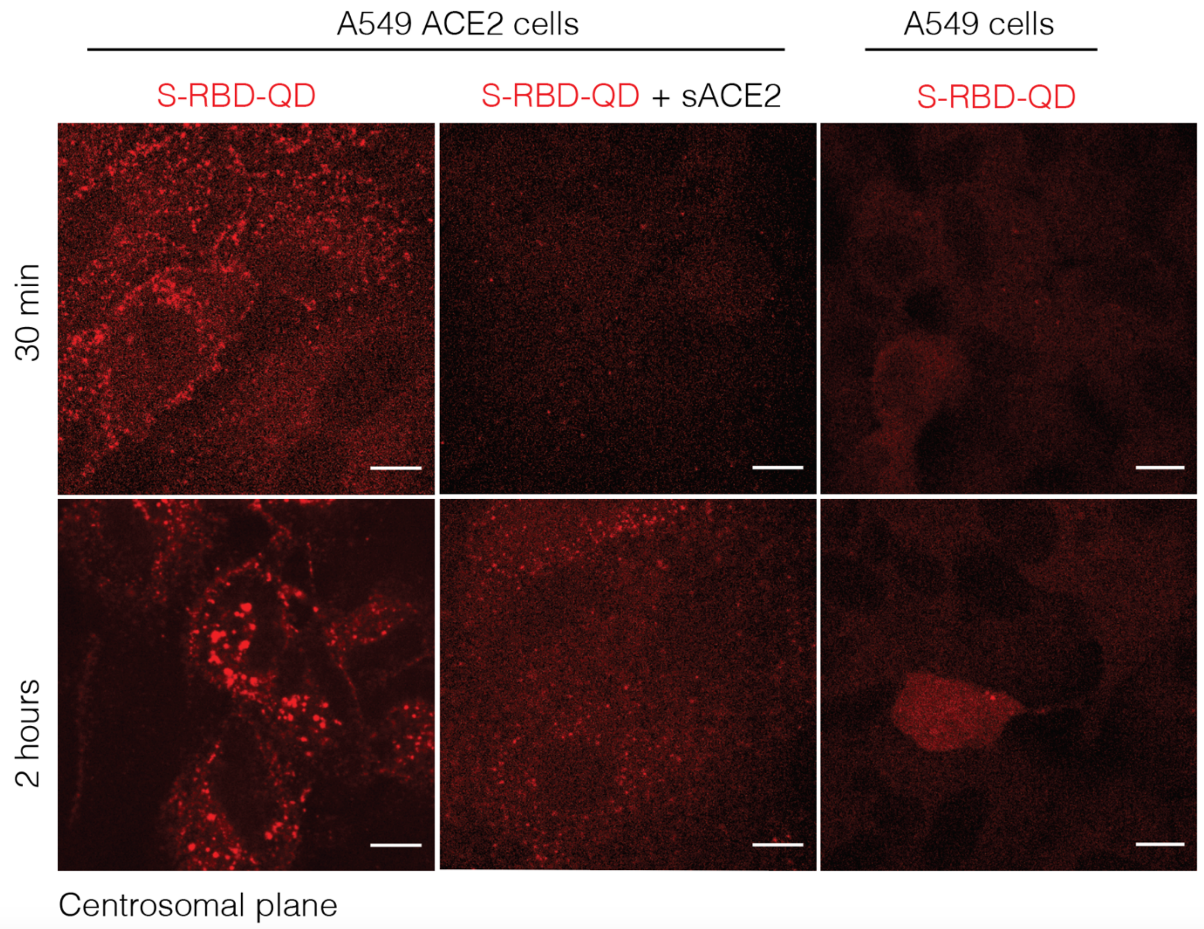


**Fig. S1. QD-Labeling target specificity controls**

Representative images of S-RBD-QD (Red) were obtained from live A549 ACE2 or A549 cells after 30 minutes and 2 hours of treatment with S-RBD (100nM) and S-RBD + sACE2. The scale bar is 10 µm.

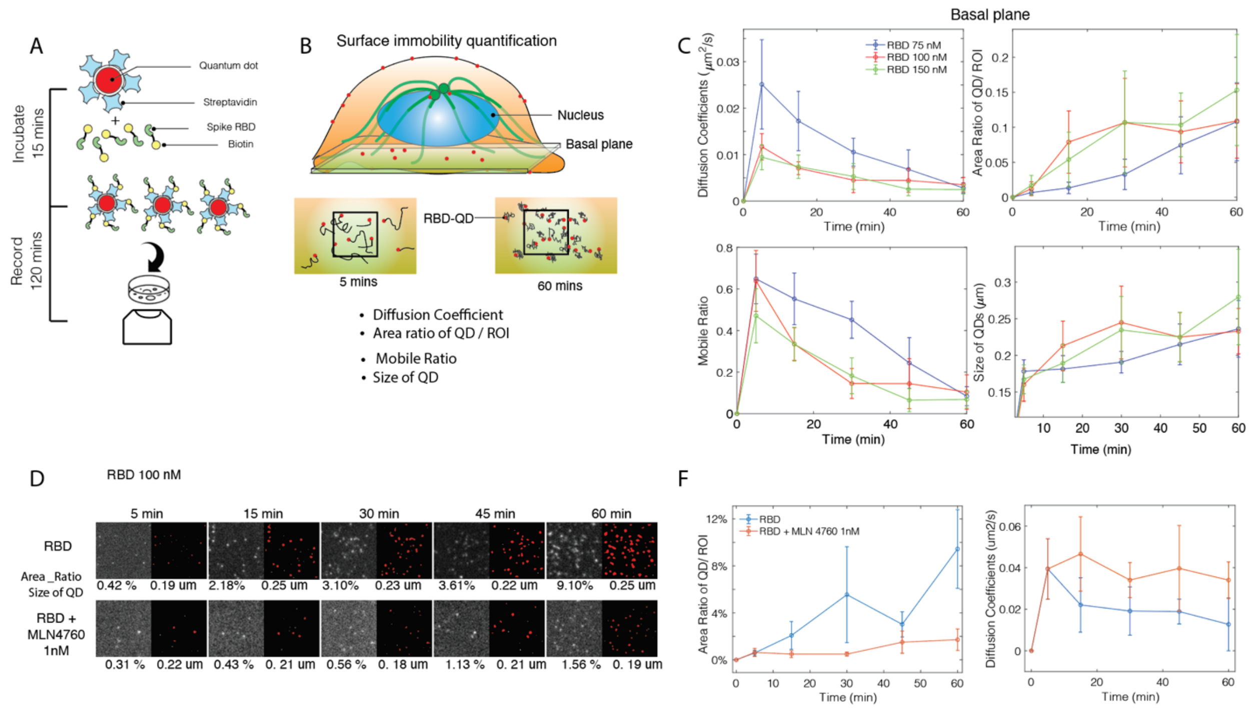


**Fig. S2. Immobility of S-RBD-QD at basal plane: quantification and ACE2 inhibitor MLN4760 treatment. (A)** Illustration of QD Labeling of ACE2 by S-RBD: S-RBD (green), biotinylated, is incubated with Quantum dot (red), conjugated with streptavidin for 15 minutes at room temperature and then applied to the live cells. Recording takes place in the following two hours. **(B)** Illustration of image capture location and quantification method of surface immobility of QDs at the ROI. To approach increasing numbers and slowing down of QDs, four quantifications: Diffusion coefficients, Area Ratio of QD vs ROI, Mobile Ratio of QDs, and the Sizes of the QD from ROI are calculated from 5 minutes to 60 minutes. **(C)** Dose dependence of QD immobility on the basal plane of A549 ACE2 cells quantified using the method introduced in (B). After the QD treatment, the basal plane videos of 2 minutes from A549 ACE2 cells are quantified at 5, 15, 30, 45, and 60 minutes for S-RBD concentrations of 75nM, 100nM, and 150nM. **(D)** Representative Images of S-RBD-QD: raw image (gray) and overlay of recognized QD area (red) in the ROI from A549 ACE2 cells treated with S-RBD and S-RBD together with ACE2 inhibitor MLN4760 (1nM) after treatment for 5, 15, 30, 45, and 60 minutes. Area Ratio of QD vs ROI, mean size of QDs is labeled under each image. **(E)** Plot of area ratio of QD vs ROI and diffusion coefficients under the conditions of RBD, and RBD + MLN4760.

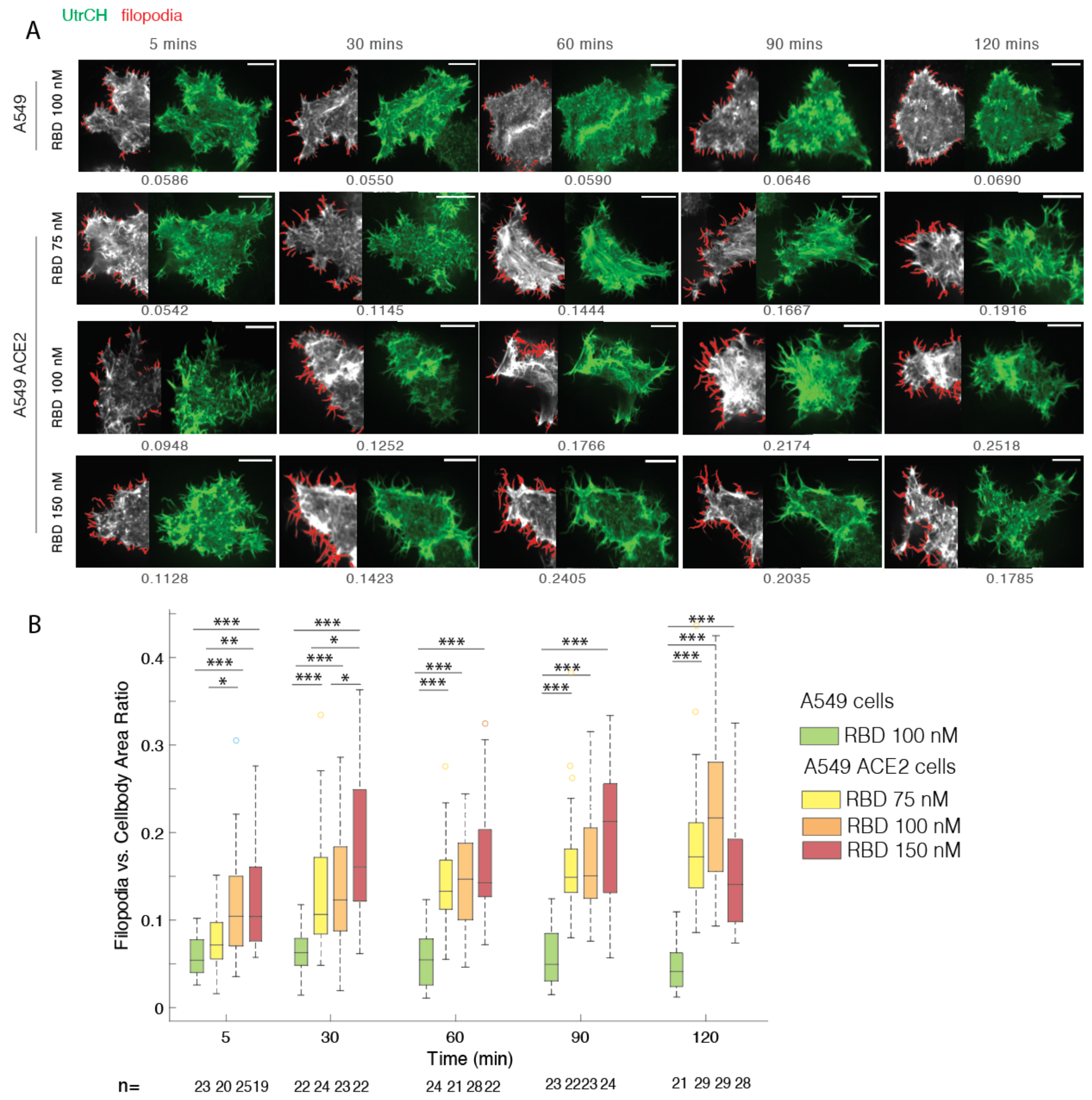


**Fig. S3. Filopodia induced by different dose of S-RBD treatment. (A)** Representative Images of Filopodia at the Basal Plane: Shown for A549 or A549 ACE2 cells, using transiently transfected UtrCH (green), at the 5, 30, 60, 90, 120 minute time points of treatment from 75nM, 100nM, and 150nM of S-RBD treatment. Scale bar is 10 µm. The left half of each image highlights the recognized filopodia area (red) overlaying with the cell (gray). The number of area ratios calculated for each example cell is presented under the image. **(B)** Filopodia vs. Cell Body Area Ratio Plots: Calculated for A549 and A549 ACE2 cells under the conditions of 75nM, 100nM, and 150nM of S-RBD treatment. (*P < 0.1,**P < 0.01; ***P < 0.001). The number of cells (n) for each calculation is presented under each box.

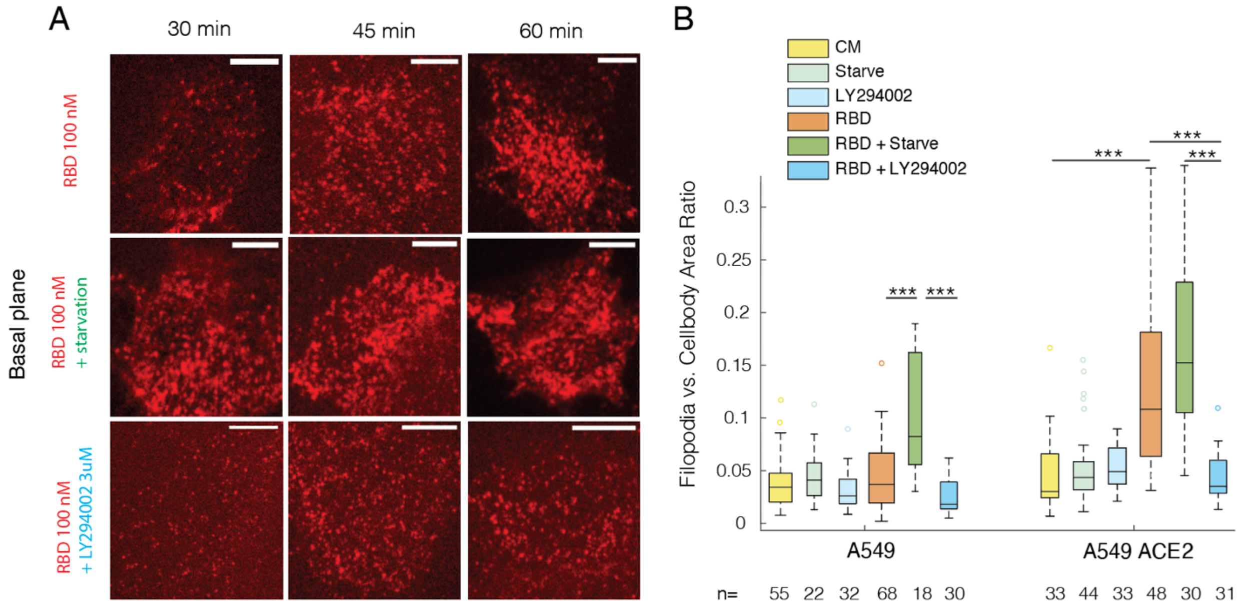


**Fig. S4. ACE2 surface immobility and filopodia in response to treatment directly targeting filopodia. (A)** Representative Time-Lapse Images of S-RBD-QD (Red): Obtained at the basal plane (as illustrated in (Fig. 1E)) from live A549 ACE2 cells after S-RBD-QD treatment for 30, 45, 60 minutes under the treatment of RBD, RBD+ starvation, and RBD + LY294002. Scale bar is 10 µm. **(B)** Filopodia vs. Cell Body Area Ratio Plots: Calculated for A549 and A549 ACE2 cells under the conditions of control, RBD, RBD+ starvation, and RBD + LY294002 treatment at the 45min time point. (***P < 0.001). The number of cells (n) for each calculation is presented under each box.

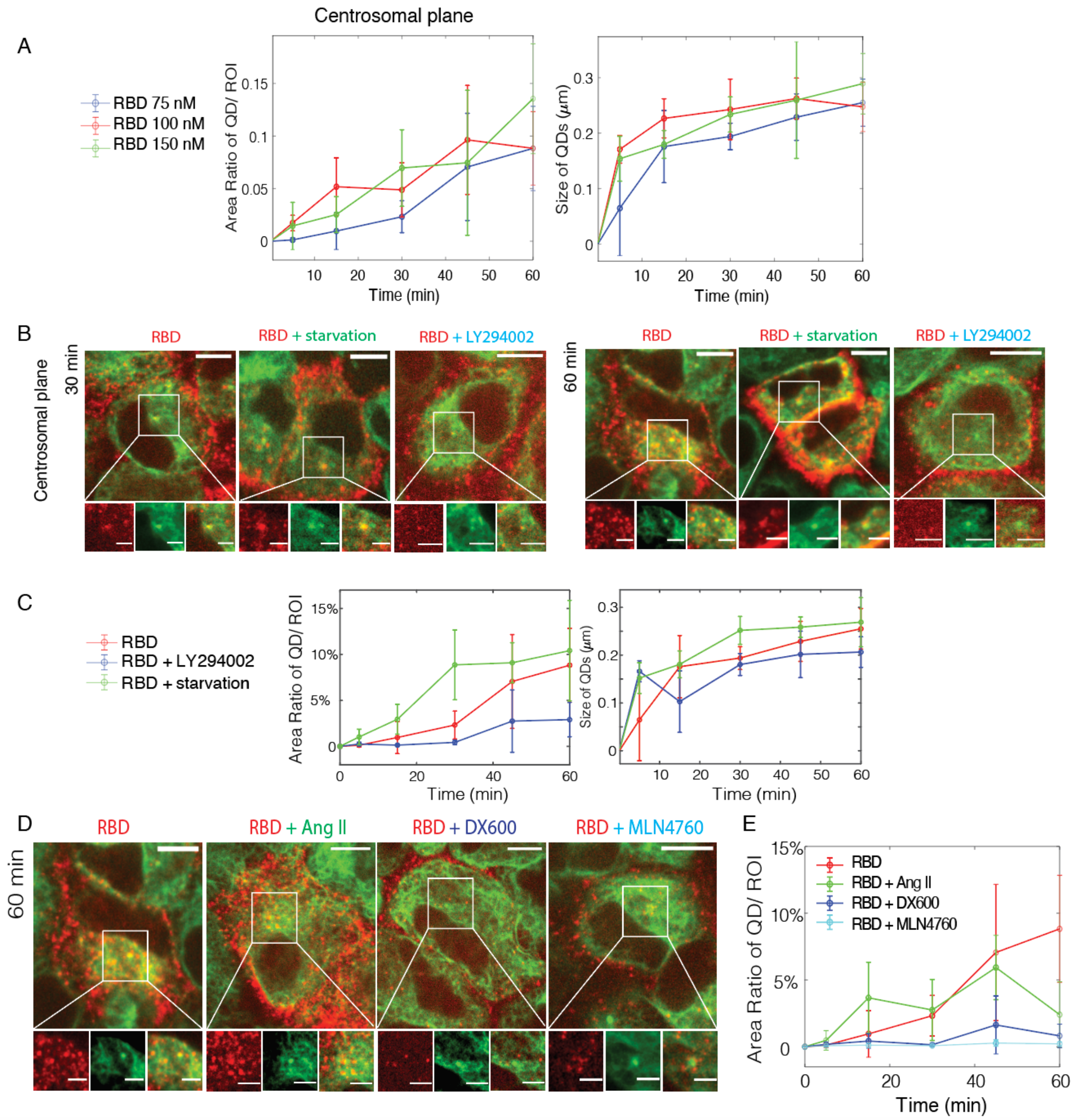


**Fig. S5. Characterization of ACE2 endocytosis labeled by S-RBD-QD. (A)** Area Ratio of QD/ROI (Red) and Size of QDs (Blue) Plots: Dose dependence of QD endocytosis on the centrosomal plane of A549 ACE2 cells to supplement data shown in Fig. 3C. After the QD treatment, the centrosomal plane videos of 2 minutes from A549 ACE2 cells are quantified at 5, 15, 30, 45, and 60 minutes for S-RBD concentrations of 75nM, 100nM, and 150nM. **(B)** Representative Images of S-RBD-QD Endosomes (Red): Shown in the whole cell view (Scale bar is 10 μm) and at the enlarged ROI selected (Scale bar is 5 μm) after 30 and 60 minutes of treatment of RBD, RBD+starvation, and RBD+ LY294002. Tubulin (green) staining shows the location of the centrosomes. (C) Area Ratio of QD/ROI (Red) and Size of QDs (Blue) Plots: the centrosomal plane videos of 2 minutes from A549 ACE2 cells are quantified at 5, 15, 30, 45, and 60 minutes for treatment of RBD, RBD+starvation, and RBD+ LY294002. (D) Representative Images of S-RBD-QD Endosomes (Red): Shown in the whole cell view (Scale bar is 10 μm) and at the enlarged ROI selected (Scale bar is 5 μm) after 60 minutes of treatment of RBD,

RBD+Ang II, RBD+ DX600 and RBD+ MLN4760. Tubulin (green) staining shows the location of the centrosomes. (E) Area Ratio of QD/ROI (Red) Plots: the centrosomal plane videos of 2 minutes from A549 ACE2 cells are quantified at 5, 15, 30, 45, and 60 minutes for treatment of RBD, RBD+Ang II, RBD+ DX600 and RBD+ MLN4760.

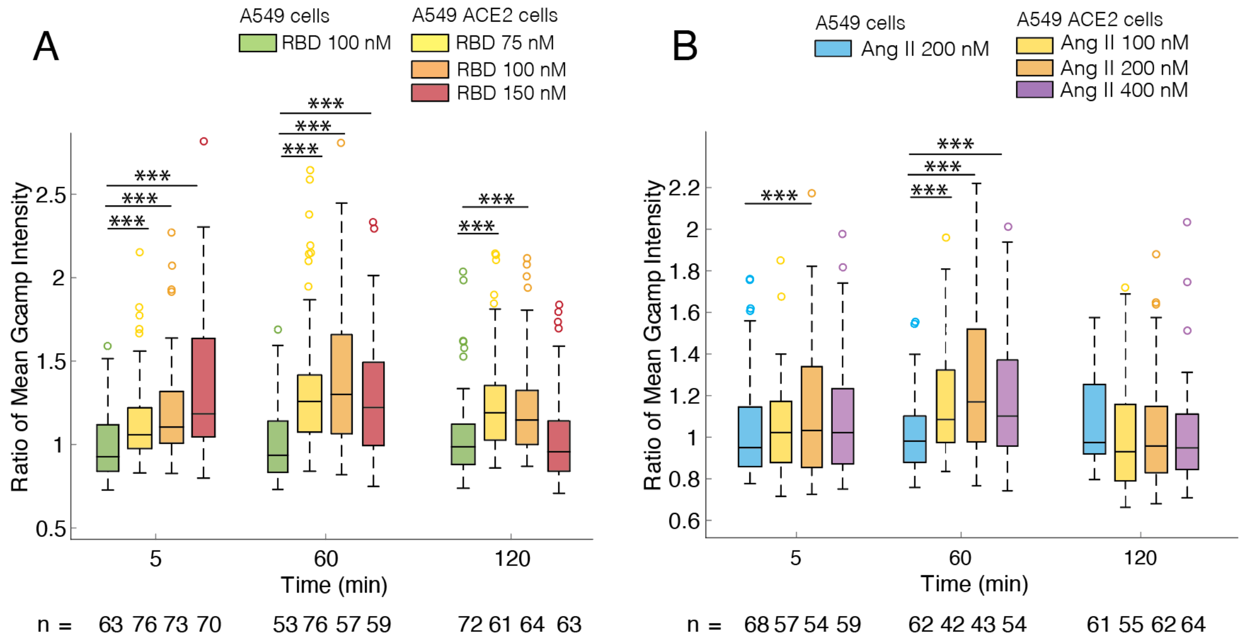


**Fig. S6. Intracellular calcium rise induced by different dose of S-RBD/Ang II treatment. (A)** Box Plots of Normalized Mean Gcamp6f Intensity: Time-lapse of each treatment was calculated for 5, 60 and 120 minutes (***$P < 0.001$). Calculated for A549 under 100nM of S-RBD and A549 ACE2 cells under the conditions of 75nM, 100nM, and 150nM of S-RBD treatment. Each measured intensity is normalized by the mean intensity of A549 ACE2 cells under control conditions. The number of cells (n) for each calculation is presented under each box. (B) Box Plots of Normalized Mean Gcamp6f Intensity: Time-lapse of each treatment was calculated for 5, 60 and 120 minutes (***$P < 0.001$). Calculated for A549 under 200nM of Ang II and A549 ACE2 cells under the conditions of 100nM, 200nM, and 400nM of Ang II treatment. Each measured intensity is normalized by the mean intensity of A549 ACE2 cells under control conditions. The number of cells (n) for each calculation is presented under each box.

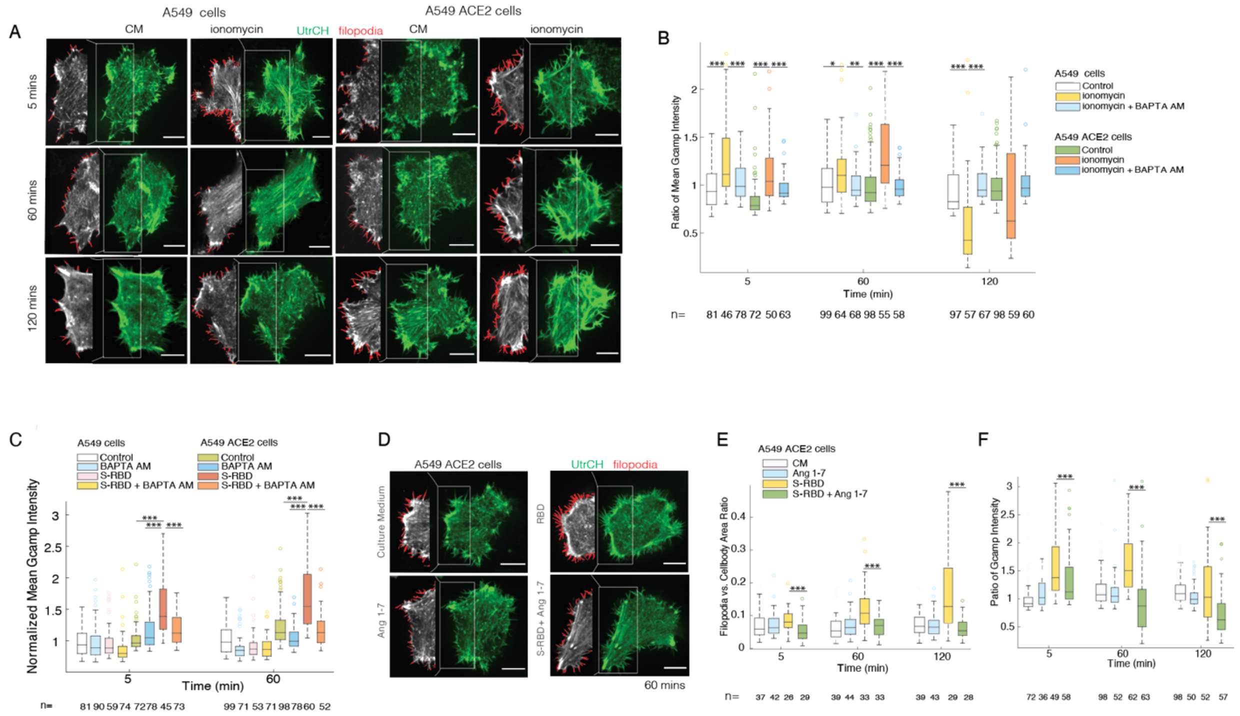


**Fig. S7. Filopodia induced by intracellular calcium rise, Ang 1-7 reverses effects on filipodia and intracellular levels caused by S-RBD ACE2 interaction. (A)** Representative Images of Filopodia at the Basal Plane: Shown for A549 or A549 ACE2 cells, using transiently transfected UtrCH (green), at the 5,60, 120-minute time point of treatment from Control and ionomycin (0.25µM) treatment. Scale bar is 10 µm. The white box highlights the recognized filopodia area (red) overlaying with the cell (gray). This image serve as supplemental to Fig. 4D. (**B, C**) Box Plots of Normalized Mean Gcamp6f Intensity: Time-lapse of each treatment was calculated for 5, 60 and 120 minutes (*P < 0.1; ***P < 0.001). Each measured intensity is normalized by the mean intensity of A549 ACE2 cells under control conditions. The number of cells (n) for each calculation is presented under each box. (B) Measured intensity of the cell body from A549 and A549 ACE2 cells under the conditions of control, ionomycin, and ionomycin+BAPTA AM treatment. (C) Measured intensity of the cell body from A549 and A549 ACE2 cells under the conditions of control, BAPTA AM, S-RBD, and S-RBD+BAPTA AM treatment. (**D**) Representative Images of Filopodia at the Basal Plane: Shown for A549 ACE2 cells, using transiently transfected UtrCH (green), at the 60-minute time point of treatment from Control, Ang 1-7,RBD, RBD+ Ang 1-7 treatment. Scale bar is 10 µm. The white box highlights the recognized filopodia area (red) overlaying with the cell (gray). (**E**) Filopodia vs. Cell Body Area Ratio Plots: Calculated for A549 ACE2 cells under the conditions of Control, Ang 1-7,RBD, and RBD+ Ang 1-7 at the 5,60, 120-minute time point of treatment (***P < 0.001). The number of cells (n) for each calculation is presented under each box. (**F**) Box Plots of Normalized Mean Gcamp6f Intensity: Time-lapse of Control, Ang 1-7,RBD, and RBD+ Ang 1-7 treatment was calculated for 5, 60 and 120 minutes (*P < 0.1; ***P < 0.001). Each measured intensity is normalized by the mean intensity of A549 ACE2 cells under control conditions. The number of cells (n) for each calculation is presented under each box.

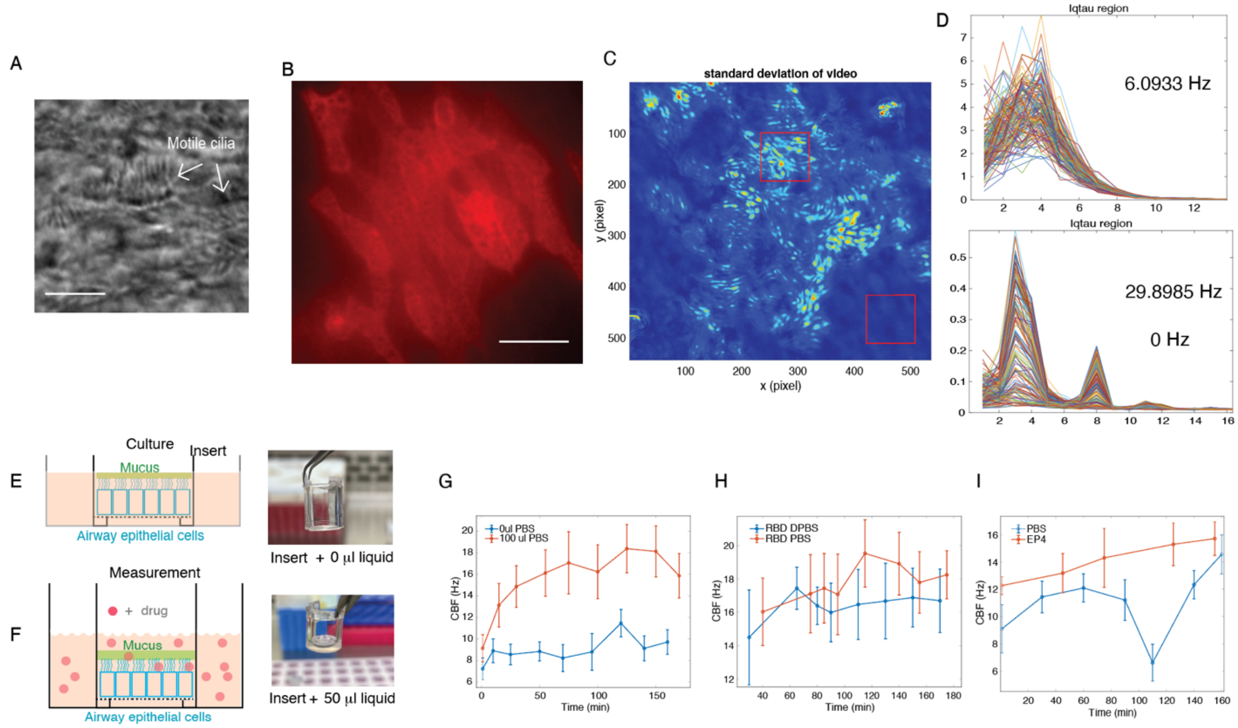


**Fig. S8. Cilia beating frequency quantification and drug treatment method demonstration.** (A) Bright field image showcase captured motile cilia. Scale bar is 10 µm. (B)

**Movie S1. Comparison of the motions of anti-ACE2-QD with and without the treatment of S-RBD.** The video depicts two example A549 ACE2 cells labeled with anti-ACE2-QD, as described in the Methods section. The scale bar for the whole-cell view is 5 µM, and for the cropped and amplified region, is 2 µM. The cell treated with S-RBD in the bottom row exhibits immobile motion, while the untreated cell in the top row displays diffusion.

**Movie S2. S-RBD-QD labeled ACE2 initiating filopodia on A549 ACE2-GFP cells.** The video showcases a representative A549 ACE2-GFP cell, along with four cropped and enlarged regions along the cell boundary, following 60 minutes of treatment with S-RBD-QD. The scale bar for the whole-cell view is 5 µM, and for the cropped and amplified region, it is 2 µM. All cropped regions highlight the de novo generation of filopodia, featuring QDs at the leading tips of growth.

**Movie S3. Motion of S-RBD-QD labeled ACE2 at UtrCH labeled filopodia on A549 ACE2 cells.** The video presents a representative A549 ACE2 cell transfected with UtrCH, accompanied by four cropped and enlarged regions along the cell boundary, following a 60-minute treatment with S-RBD-QD. The scale bar for the whole-cell view is 5 µM, and for the cropped and amplified region, it is 2 μM. All cropped regions highlight the colocalization of QD with filopodia, featuring QDs at the growing tips or moving along the filopodia.

**Movie S4. Trajectory demo from S-RBD-QD labeled ACE2 endosomes in A549 ACE2 GFP cells.** The video presents a representative A549 cell stably expressing ACE2-GFP following a 50-minute treatment with S-RBD-QD. The cell is the same as the example cell in Fig. 3D, and the trajectories are the same as shown in Fig. 3E. The pseudocolored overlay trajectories of endosomes show retrograde (orange) or antegrade transport (blue) from the centrosomes.

**Movie S5. Trajectory demo from S-RBD-QD labeled ACE2 endosomes in A549 ACE2 H374N, H378N GFP cells.** The video presents a representative A549 cell stably expressing ACE2 H374N, H378N -GFP following a 50-minute treatment with S-RBD-QD. The cell is the same as the example cell in Fig. 3D, and the trajectories are the same as shown in Fig. 3E. The pseudocolored overlay trajectories of endosomes show retrograde (orange) or antegrade transport.

**Movie S6: Bright-field capture of cilia beating motions.** The video showcases two representative regions of HNE cell cultures before and after treatment with 100 μl PBS + S-RBD for two hours. The visually detectable increase in beating frequency induced by S-RBD treatment is evident when the two conditions are compared side by side.
